# ChromSkills enables interpretable and reproducible agentic chromatin data analysis

**DOI:** 10.64898/2026.02.04.703888

**Authors:** Yuxuan Zhang, Yiman Wang, Yang Tan, Yong Zhang

**Affiliations:** State Key Laboratory of Cardiovascular Diseases and Medical Innovation Center, Institute for Regenerative Medicine, Department of Neurosurgery, Shanghai East Hospital, Shanghai Key Laboratory of Signaling and Disease Research, Frontier Science Center for Stem Cell Research, School of Life Sciences and Technology, Tongji University, Shanghai, 200092, China

## Abstract

High-throughput assays generate diverse chromatin datasets that require flexible workflows and context-dependent parameter choices. Although large language models (LLMs) can assist analysis, unconstrained LLM-based execution often exhibits unstable behavior and limited reproducibility. We present ChromSkills, a curated library of domain-specific analytical Skills for agentic chromatin data analysis on the Claude Code platform. ChromSkills encodes expert-informed decision logic and parameter-selection rules as modular, human-readable Skills and couples them with structured tool interfaces to enable reproducible execution and interpretable workflow composition from natural-language task descriptions. Across representative chromatin analysis tasks, ChromSkills consistently selected appropriate tools and parameters across repeated runs and improved execution stability and token efficiency when using Model Context Protocol-based tools. Together, ChromSkills provides a practical framework for trustworthy AI-enabled, agentic chromatin data analysis by guiding LLMs with transparent, reusable domain knowledge.

## Background

The decreasing cost of sequencing and increasing multiplexing capacity have made it routine to generate diverse chromatin datasets, including ChIP-seq, ATAC-seq, WGBS, and Hi-C, increasing the need for scalable and adaptable analysis workflows. Exploratory studies often require iterative adjustments to preprocessing steps, parameter settings, and quality-control criteria, which are difficult to accommodate with rigid, fixed pipelines. Workflow frameworks such as Nextflow [1] and Snakemake [2], and platforms such as Galaxy [3], partially address this issue by enabling custom workflow construction and community reuse. However, these systems present substantial barriers for users without strong computational training: Nextflow and Snakemake typically require scripting and familiarity with workflow configuration, while Galaxy requires users to navigate tool selection, parameterization, and data organization within a web-based environment. Higher-level, domain-specific workflow suites such as snakePipes [4] further reduce operational complexity, yet still require users to specify parameter settings and sample metadata, implicitly assuming considerable domain expertise. Moreover, in many workflow systems, analytical logic is encoded in low-level scripts or configuration files that are optimized for execution rather than interpretability, making it hard to understand the rationale for specific steps or parameter choices, inspect intermediate outputs, or troubleshoot when a workflow fails.

Large language models (LLMs) provide an opportunity to develop more intelligent and adaptive analytical workflows, given their capabilities in reasoning, modeling biological contexts, and planning tool-augmented actions [5,6]. Recent progress has moved beyond standalone models toward LLM-based agents that iteratively perceive, plan, and act, enabling dynamic adaptation to tasks with reduced human intervention. Early work has demonstrated core agent capabilities in biology, such as GeneGPT [7], which integrates LLM reasoning with external, web-based tool. Building on this, more advanced frameworks, including multi-agent systems [8,9] and instruction-tuned approaches [10,11], combine domain knowledge with iterative tool use to carry out specific biological analyses by repeatedly integrating model reasoning with external data sources and computational tools. These developments point to a progression toward increasingly autonomous and context-aware agentic systems.

Nevertheless, for chromatin data analysis, the key challenge shifts from agent design to stable and context-aware decision-making. A wide range of mature tools already exists for chromatin data analysis, such as MACS2 [12], HOMER [13], and HiCExplorer [14]. These tools provide comprehensive command-line interfaces and can, in principle, be executed automatically by an LLM-based coding agent. However, effective chromatin data analysis requires more than running individual tools. Selecting appropriate tool modes, configuring parameters, and choosing execution paths depend critically on assay type, experimental design, data quality, and the biological question. While an agent may be capable of issuing syntactically valid commands, it often lacks the domain knowledge needed to make consistent and biologically appropriate choices across heterogeneous datasets. Without explicit constraints, such agents may behave inconsistently, for example, using inappropriate parameter settings or switching tool usage across similar tasks [7]. Addressing this challenge therefore requires mechanisms for embedding expert-informed decision logic that support stable, reproducible, and context-aware analytical choices, rather than simply improving command execution.

Claude Code, a coding agent that supports extensible Skills, provides a suitable platform to address these challenges. Skills are self-contained bundles of instructions, scripts, and resources that Claude Code loads to execute specialized tasks in a repeatable manner. This design makes Skills well suited for chromatin data analysis, where stable, expert-informed decisions are required across heterogeneous assays and datasets. Here, we present ChromSkills, a curated library of modular, tool-aware Skills for chromatin data analysis. ChromSkills encodes expert decision logic as explicit analytical constraints that guide agent planning and execution, while invoking established bioinformatics tools through structured interfaces. By externalizing domain knowledge into reusable skills, ChromSkills separates analytical intent from low-level execution details, improving interpretability and reducing reliance on implicit expertise. Compared with conventional workflow systems, ChromSkills emphasizes transparent decision-making over static execution graphs, enabling adaptive yet reproducible workflows. We describe the design and implementation of ChromSkills, evaluate its stability by measuring the consistency of tool and parameter selection across repeated runs under identical conditions, and benchmark its performance against bioinformatics trainees on representative end-to-end chromatin data analysis tasks.

## Results

### Architecture of ChromSkills for agentic chromatin data analysis

ChromSkills is a curated library of domain-specific analytical Skills designed to support agentic chromatin data analysis on the Claude Code platform. The library comprises modular Skills spanning core chromatin assays, including ChIP-seq, ATAC-seq, WGBS, Hi-C, as well as integrative multi-omics analyses (Additional file 1: Fig. S1A). Rather than encoding complete analyses as fixed pipelines, ChromSkills represents chromatin data analysis as the composition of reusable analytical units, each encapsulated as an individual Skill. When a user submits a high-level analytical request in natural language, Claude Code interprets the user’s intent and constructs an analysis plan by selecting and sequencing the relevant Skills (Fig. 1A). Each selected Skill provides structured, tool-aware instructions that guide execution while enforcing assay- and context-specific constraints. If required inputs or analytical conditions are missing or ambiguous, Claude Code proactively requests clarification from the user before proceeding, preventing invalid execution paths and ensuring correct parameterization. This closed-loop interaction between user intent, Skill-guided planning, and tool-based execution enables agent-driven construction of complete chromatin data analysis workflows without manual specification of pipeline structure or tool invocation.

**Figure 1.**
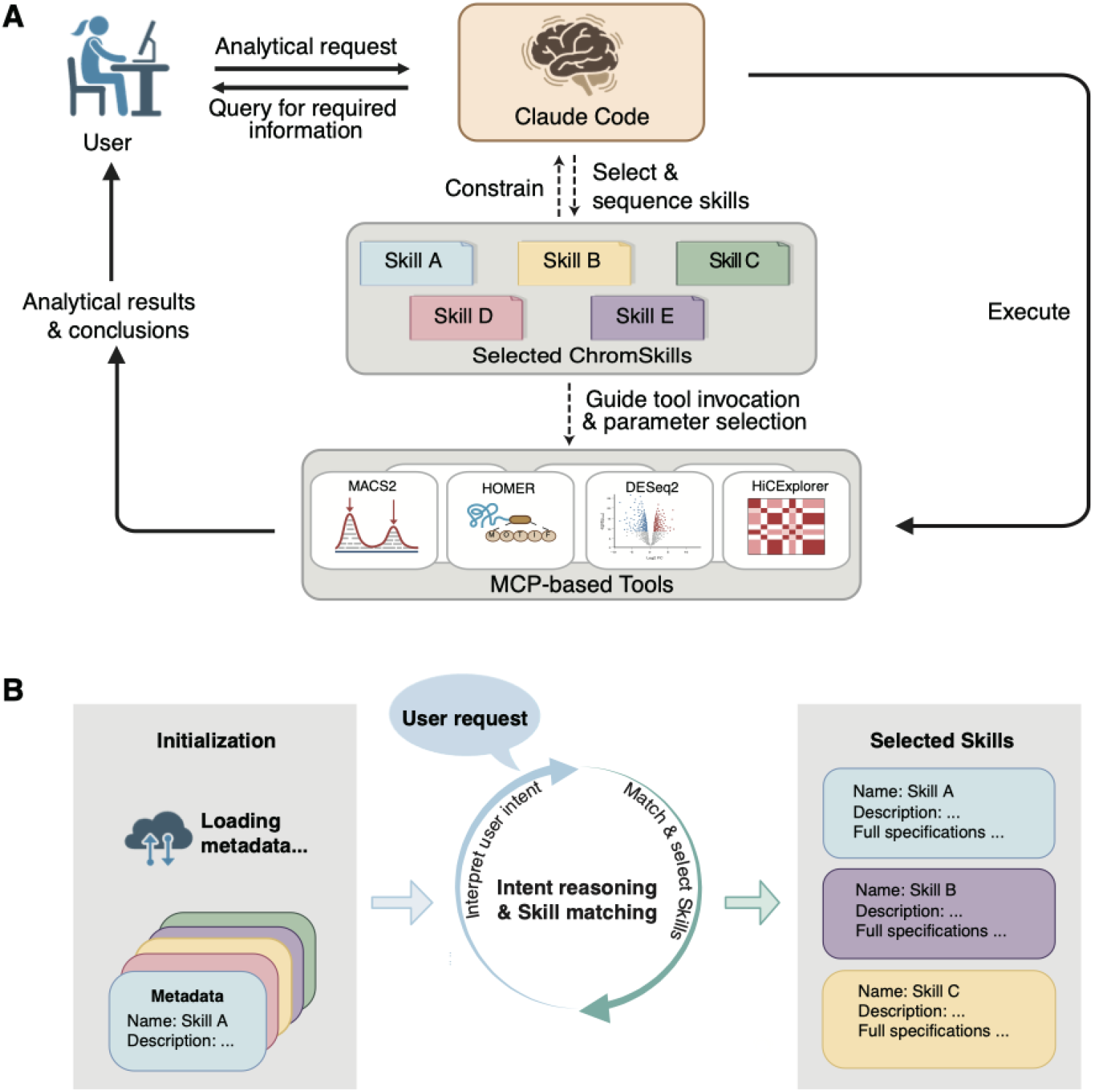
Architecture and workflow of ChromSkills. **A** Overview of the ChromSkills-enabled analytical workflow. Users provide high-level chromatin analysis requests in natural language, which are interpreted by Claude Code. The agent constructs an analysis plan by selecting and sequencing relevant ChromSkills. Skills guide execution through structured tool invocations, while missing or ambiguous inputs trigger clarification requests before execution proceeds. **B** Skill discovery and selection mechanism. Each Skill is associated with lightweight metadata (name and description) that is loaded at startup and incorporated into the system prompt. Full Skill specifications are loaded only when matched to a user request.

Skill discovery and selection are mediated through lightweight metadata associated with each Skill, including its name and a concise description (Fig. 1B). Claude Code loads this metadata at startup and incorporates it into the system prompt, enabling it to identify which Skills are relevant to a given analytical request. The full content of a Skill is loaded only when the request matches its description, allowing efficient and scalable Skill selection as the library grows. We observed that chromatin data analysis workflows are composed of a limited set of recurring analytical units that appear across diverse assays and study designs, such as alignment quality control, peak calling, genomic feature annotation, and differential-region analysis (Additional file 1: Fig. S1B). ChromSkills explicitly decomposes workflows into these reusable units, each implemented as an independent Skill. As a result, the same Skill can be reused across multiple analytical contexts without requiring users to explicitly specify which tools or workflow components to invoke. Together, these design choices establish ChromSkills as a modular framework for translating high-level analytical intent into structured chromatin data analysis workflows through principled recombination of expert-curated Skills.

### Structured Skill specifications make analytical logic explicit

To make agent-driven chromatin data analysis transparent and interpretable, each Skill in ChromSkills is defined as a structured, self-contained specification composed of three human-readable components: Overview, Inputs & Outputs, and Decision Tree (Fig. 2). This structure is designed to expose analytical intent and decision logic explicitly, rather than leaving these choices implicit in scripts or configuration files.

**Figure 2.**
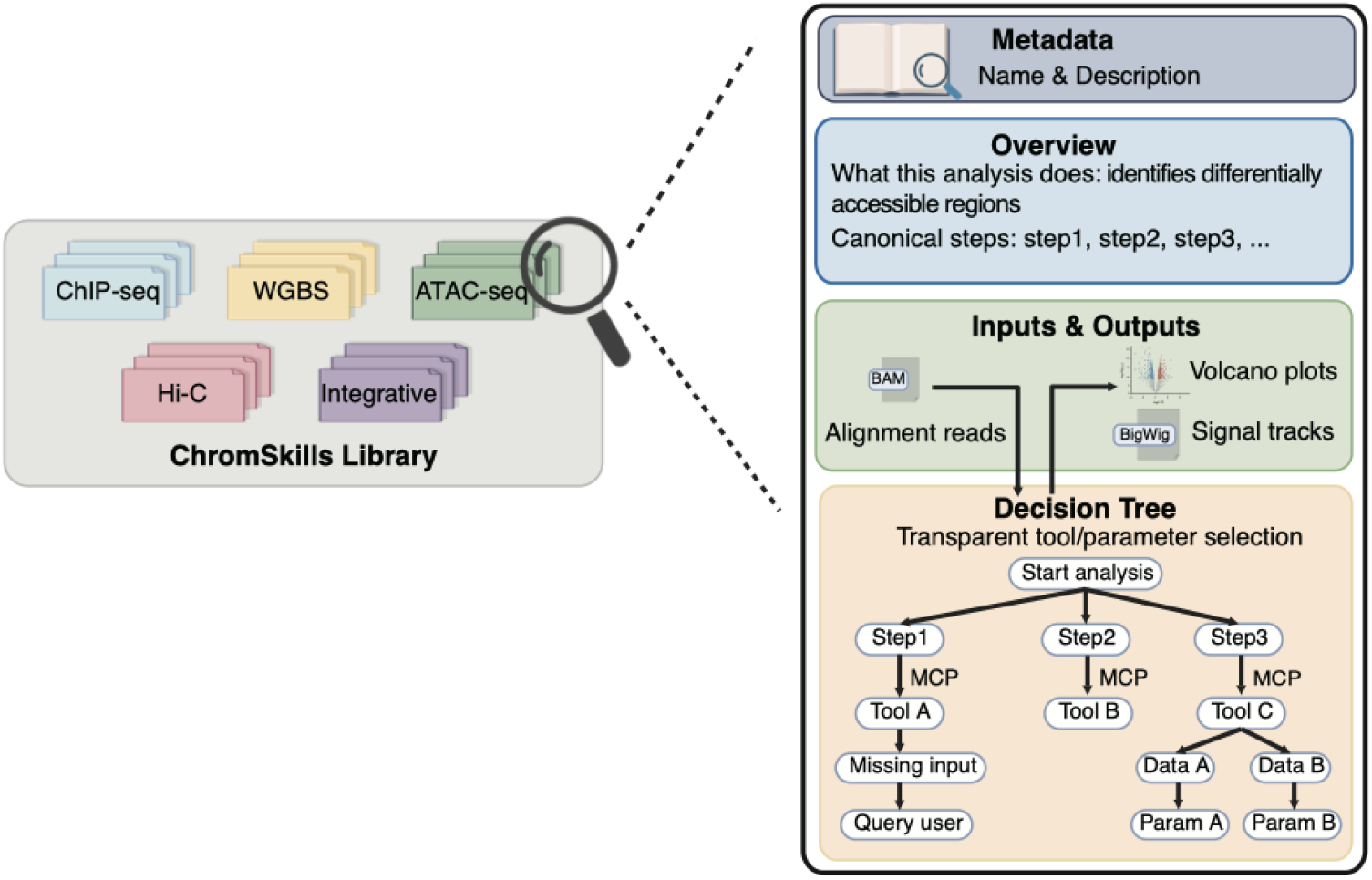
Schematic representation of the internal structure of a ChromSkill. Each Skill is defined as a self-contained, human-readable specification consisting of three components: Overview, describing the analytical goal and canonical steps; Inputs & Outputs, defining required data types, formats, and standardized outputs; and Decision Tree, encoding domain-specific conditional rules that govern tool selection, parameter choices, and execution order.

The Overview section describes the conceptual purpose of the Skill and outlines the canonical analytical steps it performs, providing a high-level procedural summary that frames subsequent actions. It also specifies the information that must be provided by the user to ensure correct execution. The Inputs & Outputs section defines required input data types, expected file formats, and standardized output artifacts, promoting consistency across Skills and facilitating interoperability between multi-step analyses. The Decision Tree section encodes domain-specific analytical logic as a sequence of conditional rules that govern tool selection, parameter determination, and execution order. Rather than expressing this logic in executable code, Decision Trees are written in natural language, making them directly inspectable by users while remaining interpretable by Claude Code. In practice, Decision Trees capture assay- and context-specific branching points such as selecting tool modes, applying dataset-dependent parameter adjustments, and choosing alternative execution paths or querying user for more information when required inputs are unavailable.

### Skill-encoded decision logic enables reproducible execution

In chromatin data analysis, tool modes and parameter settings often depend on assay type, experimental design, and data characteristics, and small deviations can lead to incorrect execution or inconsistent outputs. To assess whether ChromSkills supports context-aware and reproducible execution, we evaluated its performance on three representative chromatin data analysis tasks spanning distinct data modalities and analytical objectives (Additional file 1: Fig. S2A). Each task was executed multiple times under identical input conditions (see Methods), and we tracked the consistency of tool selection, parameterization, and overall execution outcomes across repetitions. Across all tasks and repeated runs, ChromSkills consistently selected appropriate tools and parameters, yielding stable and reproducible execution behavior (Fig. 3A). This consistency is driven by Skill-encoded decision logic that constrains tool modes and parameter choices in a context-dependent manner, rather than relying on free-form command generation, thereby reducing run-to-run variability and limiting failures caused by inconsistent parameterization.

**Figure 3.**
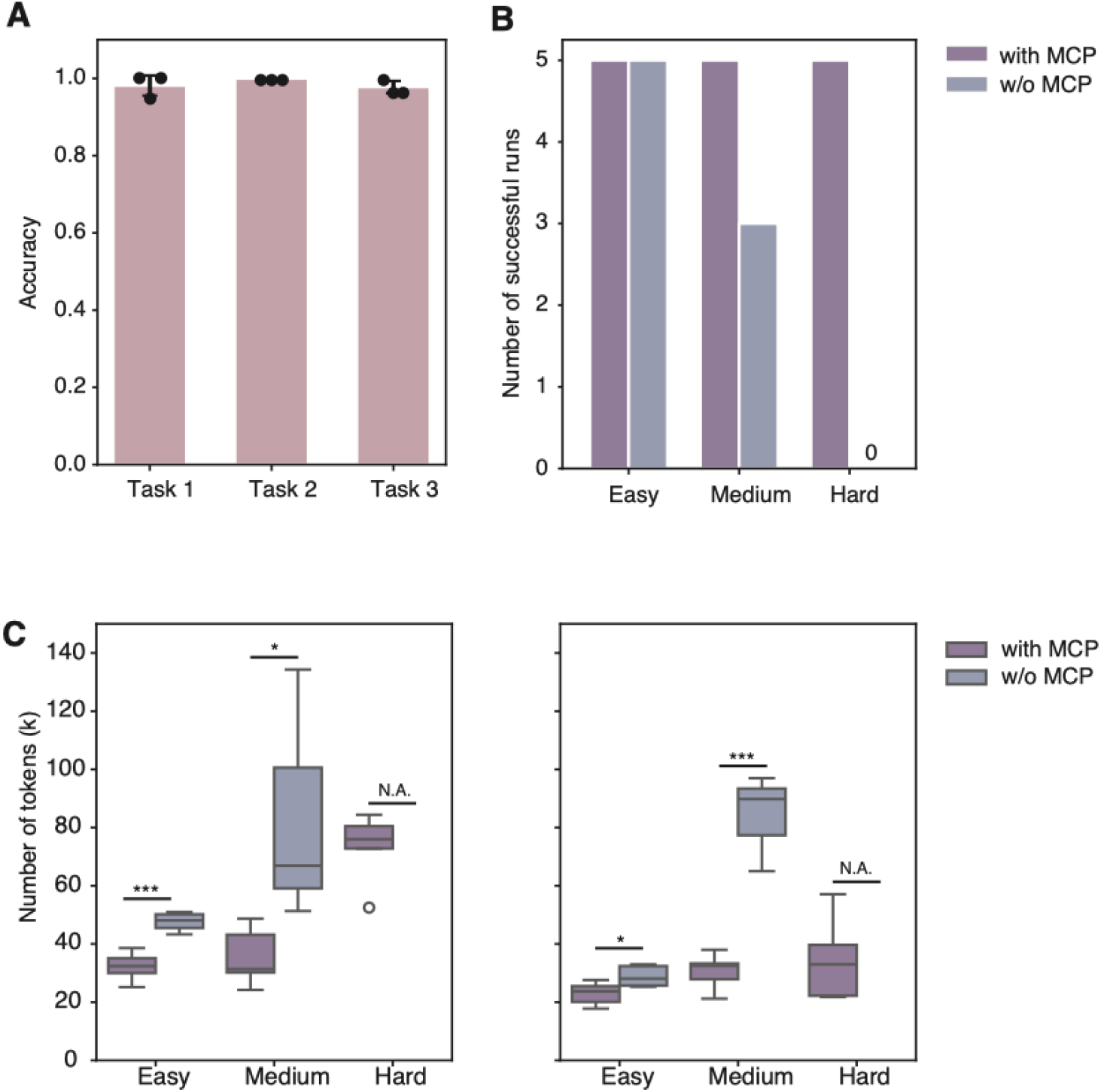
Evaluation results of execution accuracy, stability and efficiency of ChromSkills. **A** Accuracy of tool and parameter selection across repeated executions of three representative chromatin analysis tasks under identical conditions. Each value on the bar is calculated by the proportion of truly selected parameters within all parameters that required to specify during task execution. **B** Number of successful task completions across three analytical tasks of increasing difficulty (easy, medium, hard), each executed five independent times with MCP-based tools enabled (dark purple) or disabled (purple). **C** Distribution of input and output token usage with MCP-based tools enabled (dark purple) or disabled (purple). Statistical significance was evaluated using a two-sided independent t-test. ***p < 0.001, *p<0.05.

In addition to Skill-level constraints, we further constrained execution by encapsulating standardized sub-workflows as Model Context Protocol (MCP)-based tools [15] and guiding the agent to select context-appropriate parameters through structured interfaces. To quantify the impact of MCP-based tool encapsulation, we used a separate set of three analytical tasks of increasing difficulty (Additional file 1: Fig. S2B) and executed each task five independent times under two settings: with MCP-based tools enabled and without MCP-based tools (see Methods). MCP-based tools increased success rates across all difficulty levels, with the largest gains observed for medium and hard tasks where failures were common without MCP (Fig. 3B). MCP-based tools also reduced both input and output token usage (Fig. 3C), consistent with reduced iterative debugging triggered by incorrect or non-executable commands. Together, these analyses indicate that reproducible agentic chromatin data analysis benefits from complementary constraints: Skill-encoded decision logic that stabilizes tool and parameter selection, and structured tool interfaces that improve execution robustness and efficiency.

### ChromSkills demonstrates practical analytical competence in real-world settings

We benchmarked the ChromSkills-enabled Claude Code system against bioinformatics trainees using three representative chromatin data analysis tasks spanning increasing complexity and including both single-modality and integrative multi-omics analyses (Fig. 4A, Additional file 1: Fig. S2A). Each task was provided to both ChromSkills and human participants, and all outputs were submitted as structured analytical reports to enable format-consistent evaluation. Human participants were undergraduate trainees who had recently completed a computational genomics course; approximately 30 trainees participated in each task. Reports were scored using a holistic framework that assessed methodological appropriateness and analytical correctness (see Methods). Each report was independently scored by both ChatGPT and the instructor of the computational genomics course. Across the three tasks, ChromSkills ranked near the top of the score distribution and outperformed approximately 80-97% of trainees. This pattern was observed in evaluations from both ChatGPT and the instructor. The largest difference between ChromSkills and the average trainee score was observed for the task involving cross-modality integration (Fig. 4B).

**Figure 4.**
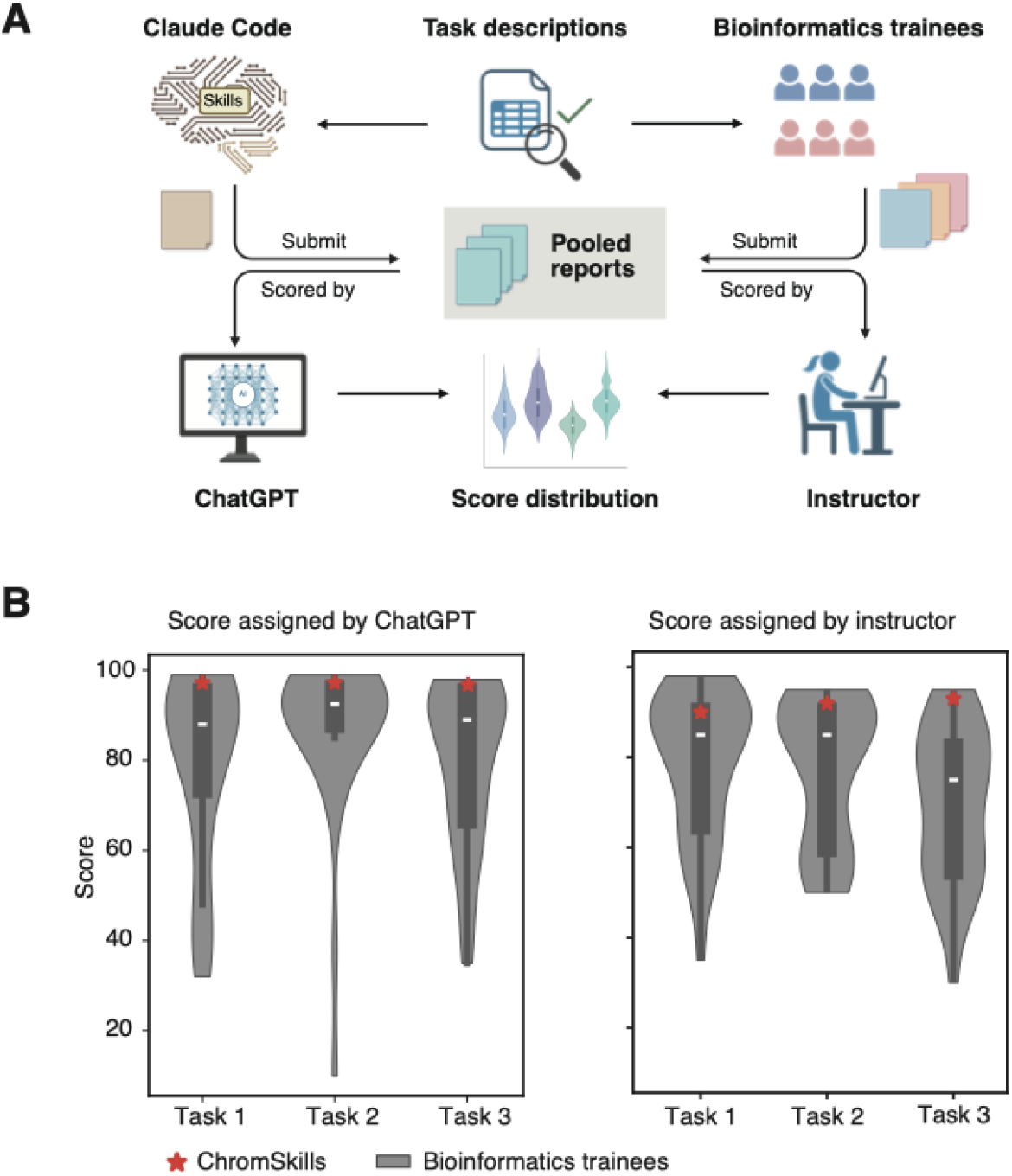
Benchmarking ChromSkills against bioinformatics trainees. **A** Benchmarking design. Three end-to-end chromatin data analysis tasks were independently completed by ChromSkills-enabled Claude Code and by bioinformatics trainees. All outputs were submitted as structured analytical reports and evaluated by both ChatGPT and the instructor of the computational genomics course. **B** Score distributions for ChromSkills and bioinformatics trainees across three chromatin data analysis tasks, as assessed by ChatGPT (left) and the instructor of the computational genomics course (right).

We next tested whether ChromSkills could reproduce representative analyses from a published ATAC-seq study [16] using only natural-language prompts, without manual pipeline construction or manual parameter tuning. The original study examined chromatin accessibility changes in myofibers across physiological and pathological conditions, including injury response and dystrophic disease models. Using ChromSkills, we requested standard ATAC-seq analyses consistent with the study design, including BAM filtration, peak calling, genomic feature annotation, transcription start site (TSS) enrichment analysis, differential accessibility analysis, and functional enrichment (Additional file 1: Fig. S3). ChromSkills assembled the workflow by selecting and ordering the corresponding Skills and executed the analyses with context-appropriate parameters. The main qualitative patterns reported in the study were reproduced (Additional file 1: Fig. S4), including genomic distributions of peaks across annotated features, enrichment of accessibility around transcription start sites, and Gene Ontology terms related to muscle function and injury response among genes associated with accessibility changes. ChromSkills also identified condition-associated regulatory regions with significant accessibility changes and summarized these results using standard visualizations and gene proximity annotations. Genes linked to condition-specific peaks showed differences in enriched functional terms between MDX and WT conditions. Together, these results show that ChromSkills can reproduce end-to-end chromatin data analysis workflows using only natural-language task descriptions.

## Discussion

ChromSkills addresses a central limitation in chromatin data analysis: the difficulty of achieving flexible automation while maintaining stability, interpretability, and reproducibility. Although many mature tools exist for chromatin data analysis, effective analysis remains constrained by manual workflow construction and context-dependent parameter selection. Our results indicate that modular, domain-informed Skills contribute to stable execution and high-quality analysis outputs across diverse tasks.

A key design principle of ChromSkills is the separation of analytical reasoning from low-level execution. By encoding expert decision logic in human-readable Skills and delegating deterministic execution to MCP-based tools, ChromSkills reduces common failure modes of LLM-based analysis, including hallucinated commands and unstable parameter choices. This design is consistent with the observed improvements in success rate and token efficiency, as well as the consistent reuse of Skills across heterogeneous workflows without explicit user instruction. Importantly, this approach preserves flexibility by enabling task-driven workflow composition rather than relying on fixed pipelines. ChromSkills is intentionally designed for lightweight, interactive analysis in local desktop or laptop environments and therefore focuses on post-alignment workflows starting from processed inputs, rather than computationally intensive preprocessing steps such as read mapping that are typically performed once using established pipelines on high-performance computing systems. ChromSkills also improves the usability of chromatin data analysis by making analytical decisions transparent. Unlike traditional workflows that embed logic in scripts or configuration files, ChromSkills exposes tool usage and parameter-selection logic in natural language through structured Skill specifications, enabling users to inspect how analyses are performed and how conditional decision rules guide tool and parameter choices.

Despite these advantages, ChromSkills also has limitations. ChromSkills currently focuses on widely used chromatin assays and established analytical paradigms, and its performance depends on the completeness and quality of encoded domain knowledge. Extending ChromSkills to emerging assays, novel experimental designs, or highly unconventional analyses will require continued curation and validation by domain experts. In addition, while Skill-level decision trees constrain parameter selection effectively, they do not eliminate the need for user judgment in ambiguous biological scenarios, such as conflicting quality metrics or unconventional experimental designs. In benchmarking against undergraduate trainees and in reproduction of a published ATAC-seq study, ChromSkills ranked near the top of the score distribution and recapitulated the main qualitative patterns from the original analysis using natural-language task descriptions alone. Future work could further strengthen robustness by expanding assay coverage, maintaining Skills as tools and best practices evolve, and providing clearer decision support when inputs or experimental context are under-specified.

## Conclusion

We present ChromSkills, a library of domain-specific Skills that enables robust, interpretable, and agentic chromatin data analysis by constraining LLM-based analysis with explicit expert knowledge. By decomposing chromatin data analysis workflows into reusable Skills and encoding analytical intent, decision logic, and parameter selection in natural language, ChromSkills improves execution stability, readability, and reproducibility while preserving analytical flexibility. Benchmarking against undergraduate bioinformatics trainees shows that ChromSkills can rank near the top of the score distribution across diverse chromatin analysis tasks, and ChromSkills reproduced the main qualitative patterns of a published ATAC-seq analysis using only natural-language task descriptions. Together, ChromSkills illustrates an approach to AI-assisted data analysis in which LLMs are guided by transparent, reusable domain knowledge rather than relying on unconstrained code generation.

## Methods

### ChromSkills implementation

ChromSkills is implemented as a collection of modular, domain-specific Skills for chromatin data analysis within the Claude Code platform. To ensure execution reproducibility and dependency isolation, Claude Code and all required bioinformatics software are deployed in a unified Docker container. ChromSkills is designed for lightweight, interactive post-alignment analysis and therefore starts from processed inputs (e.g., BAM files) rather than including computationally intensive steps such as read mapping. Claude Code provides the Skill-loading mechanism and orchestrates tool invocation during analysis, while the LLM used for reasoning and code generation in this work is DeepSeek V3.2 [17]. Skills are discovered and loaded according to Claude Code’s directory and formatting conventions; therefore, all ChromSkills follow the platform’s native Skill specification. Each Skill is implemented as a Markdown document that encodes the Skill’s instructions and decision logic in a human-readable format.

Executable components required by ChromSkills are provided through MCP-based tools implemented in Python using the FastMCP framework [18]. These tools expose structured, parameterized interfaces to the underlying bioinformatics software within the container. When an analysis step requires external computation, the corresponding Skill instructs Claude Code to invoke a specific MCP tool by name and to provide parameters determined by the Skill’s decision logic (see Code availability).

### Evaluation of parameter selection accuracy under ChromSkills decision rules

To evaluate the consistency and appropriateness of automatic parameter selection, we assessed ChromSkills on three representative chromatin data analysis tasks spanning different data modalities and analytical objectives, including both single-modality and integrative multi-omics workflows (Additional file 1: Fig. S2A). Each task was executed independently three times under identical conditions, using the same input datasets and the same task prompt for all repetitions.

Prior to execution, we predefined the set of context-dependent parameters that required explicit specification during each workflow (Additional file 2: Table S1), including tool variants, normalization strategies, assay-specific flags, and key threshold values. For each run, we recorded the parameters values selected during execution and compared them to the corresponding reference choices specified by the expert-curated decision logic encoded in the relevant ChromSkills. Parameter selection accuracy for a given run was quantified as the proportion of parameters whose selected values matched the reference choices defined by the Skill-level decision trees.

### Evaluation of MCP-based tools for execution stability and efficiency

To quantify the contribution of MCP-based tools to execution stability and computational efficiency, we designed three representative chromatin data analysis tasks of increasing difficulty (Additional file 1: Fig. S2B). Each task was executed independently five times under two settings: (i) with MCP-based tools enabled and (ii) with MCP-based tools disabled. In the MCP-disabled setting, Skill instructions guided the agent to follow the same analytical logic and sub-workflow structure implemented in the corresponding MCP tools, but required the agent to generate and execute the underlying command-line steps directly.

Task completion was evaluated using a predefined success criterion. A run was considered successful if all required outputs were generated and the workflow progressed through all steps without incorrect tool usage and with no more than three debugging attempts. A run was classified as a failure if the required outputs were not produced, or if more than three execution errors or debugging attempts occurred. Execution errors included syntactically invalid commands, incorrect tool usage, and runtime failures that prevented progression to subsequent steps.

### Benchmarking ChromSkills performance against trainees

To benchmark the analytical performance of ChromSkills against human practitioners, we compared the ChromSkills-enabled Claude Code system with undergraduate bioinformatics trainees on three end-to-end chromatin data analysis tasks (Additional file 1: Fig. S2A). Each task was administered to both ChromSkills and human participants using identical task descriptions and the same required input data types. All participants submitted structured analytical reports describing the analysis workflow, key parameter choices, and resulting outputs; command-level details were included when available. For ChromSkills, no external assistance was provided during task completion other than responses to clarification questions explicitly issued by Claude Code; tool selection, parameterization, and execution were performed within the ChromSkills framework.

All submitted reports generated by either ChromSkills or human participants were evaluated using the same predefined, holistic scoring framework (Additional file 3: Table S2). The rubric assessed methodological appropriateness and analytical correctness to enable direct comparison across human and ChromSkills-generated analyses. Each report was independently scored by two evaluators (ChatGPT and the course instructor), and benchmark outcomes were summarized by comparing task-specific score distributions and reporting the percentile rank of ChromSkills relative to the trainee distribution.

### Reproduction of a published ATAC-seq study

To evaluate ChromSkills on a realistic public dataset, we reproduced representative analyses from a published ATAC-seq study [16]. Raw sequencing data were downloaded from the Gene Expression Omnibus (GEO). Read alignment was performed outside ChromSkills as a one-time preprocessing step: reads were aligned to the mouse genome (mm10) using Bowtie2 [19] with standard ATAC-seq-compatible parameters. The resulting BAM files were then used as the inputs to ChromSkills for all downstream analyses. Starting from the aligned BAM files, downstream analyses were performed using ChromSkills based on high-level natural-language prompts describing the analytical objectives (Additional file 1: Fig. S3). ChromSkills automatically selected and assembled the relevant Skills to construct complete ATAC-seq analysis workflows.

## Supporting information

Addtional file 1

Addtional file 2

Addtional file 3

## Data availability

The ATAC-seq dataset used for the reproduction analysis is publicly available from GEO under accession number GSE173676.

## Code availability

The source codes of ChromSkills are available on the GitHub repository (https://github.com/TongjiZhanglab/ChromSkills).

## Acknowledgments

We thank Dr. Qi Wang, the instructor of the computational genomics course, for coordinating the benchmarking exercise and independently evaluating submitted reports. We also thank the students in the course for their participation and for providing the trainee reports used in this study. This work was supported by the National Natural Science Foundation of China (32325012, 32488101), the National Key Research and Development Program of China (2021YFA1302500), and the Science and Technology Commission of Shanghai Municipality (23JS1401200).

## Author contributions

Y(Yong).Z. conceived the research, Y(Yuxuan).Z. and Y.W. developed the ChromSkills, Y(Yuxuan).Z and Y.T. tested the stability of ChromSkills, Y(Yuxuan).Z. and Y(Yong).Z. wrote the manuscript.

