## Supplementary material for "ChromSkills enables interpretable and reproducible agentic chromatin data analysis": Addtional file 1

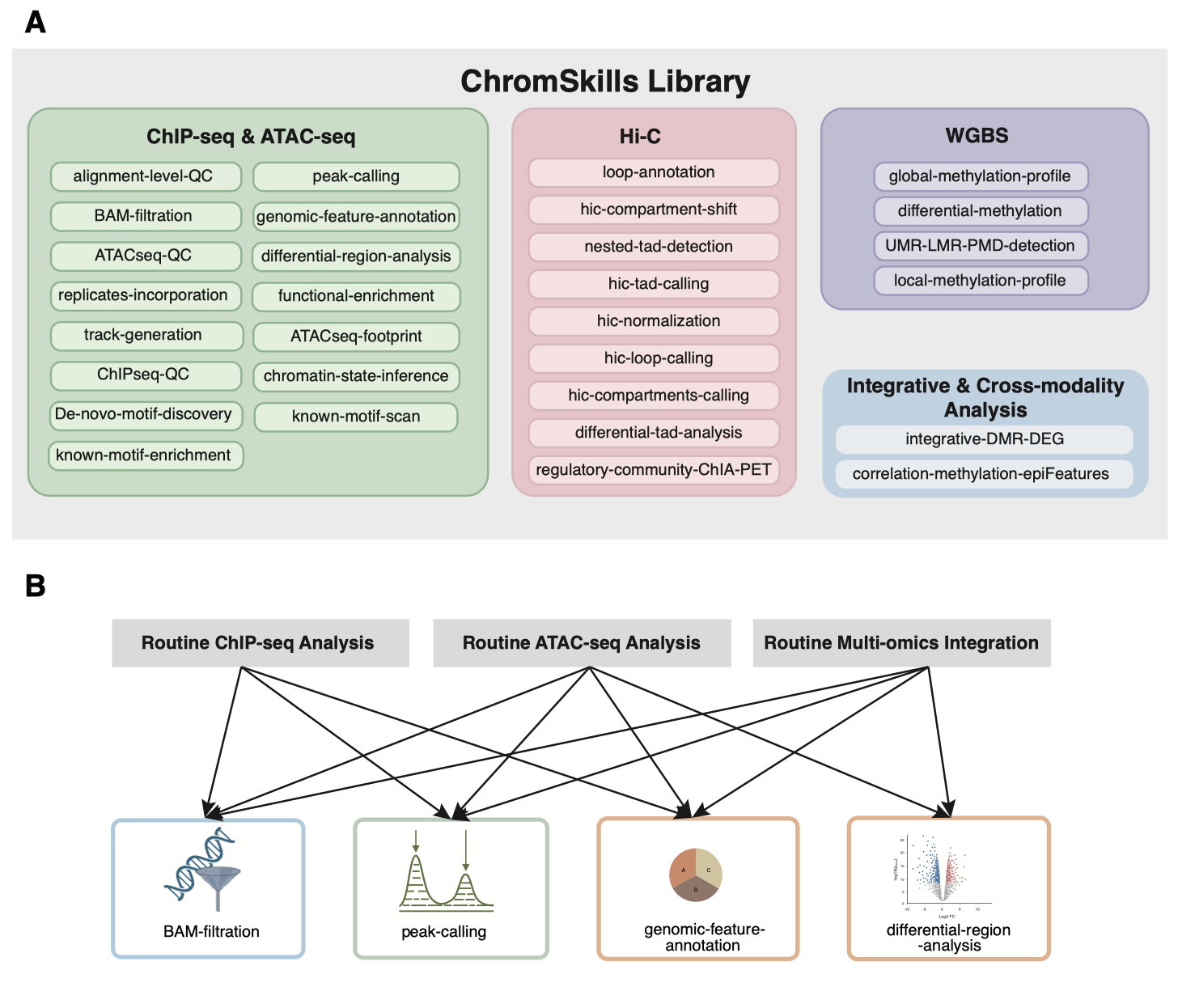

**Figure S1.** Scope and modular organization of the ChromSkills library. **A** Coverage of ChromSkills across major chromatin assays, including ChIP-seq, ATAC-seq, WGBS, Hi-C, and integrative multi-omics analyses. **B** Analytical units appear cross three routine chromatin data analyses, such as quality control, peak calling, genomic annotation, and differential-region analysis.

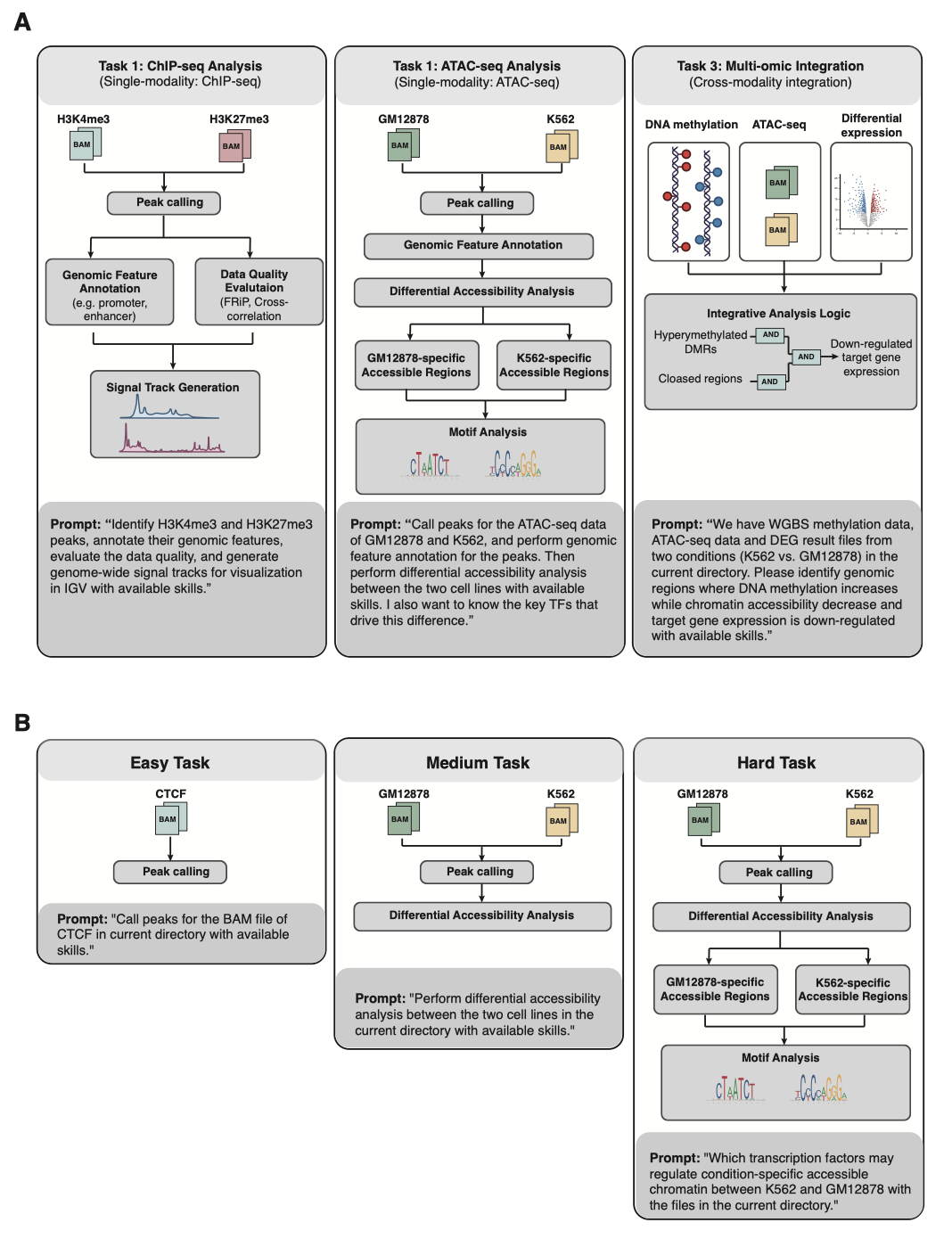

**Figure S2.** Design of evaluation tasks. **A** Three representative chromatin data analysis tasks used to assess reproducibility and benchmark ChromSkills against trainees. **B** Three additional tasks of increasing difficulty (easy, medium, hard) used to evaluate execution stability and efficiency with and without MCP-based tools.

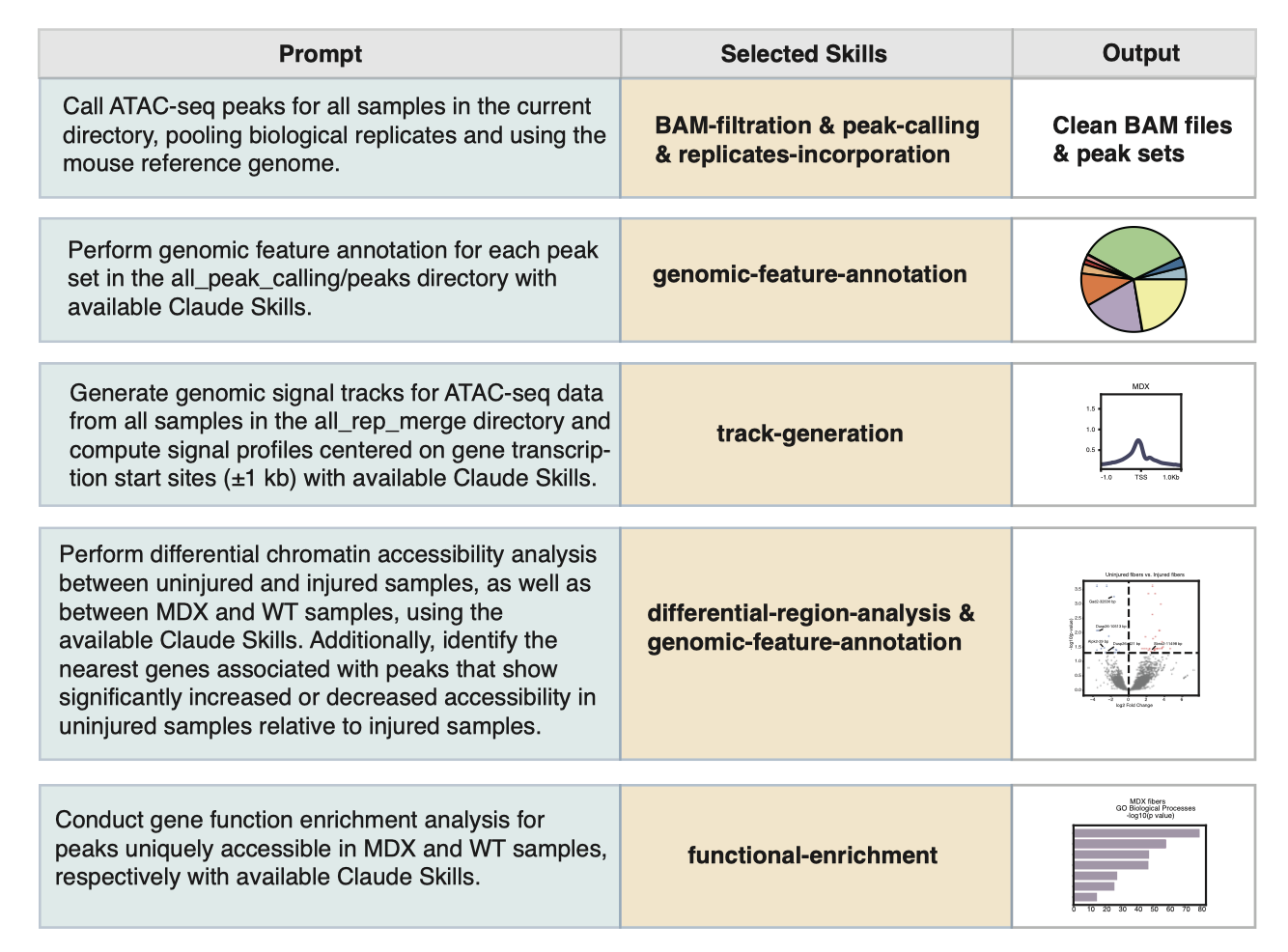

**Figure S3.** Prompts and Skills used for the reproduction of a published ATAC-seq analysis. Workflow assembled by ChromSkills in response to high-level natural-language prompts requesting standard ATAC-seq analyses, including BAM filtration, peak calling, genomic feature annotation, TSS enrichment analysis, differential accessibility analysis, and functional enrichment. Skills used in each step were selected and sequenced by ChromSkills-enabled Claude code automatically.

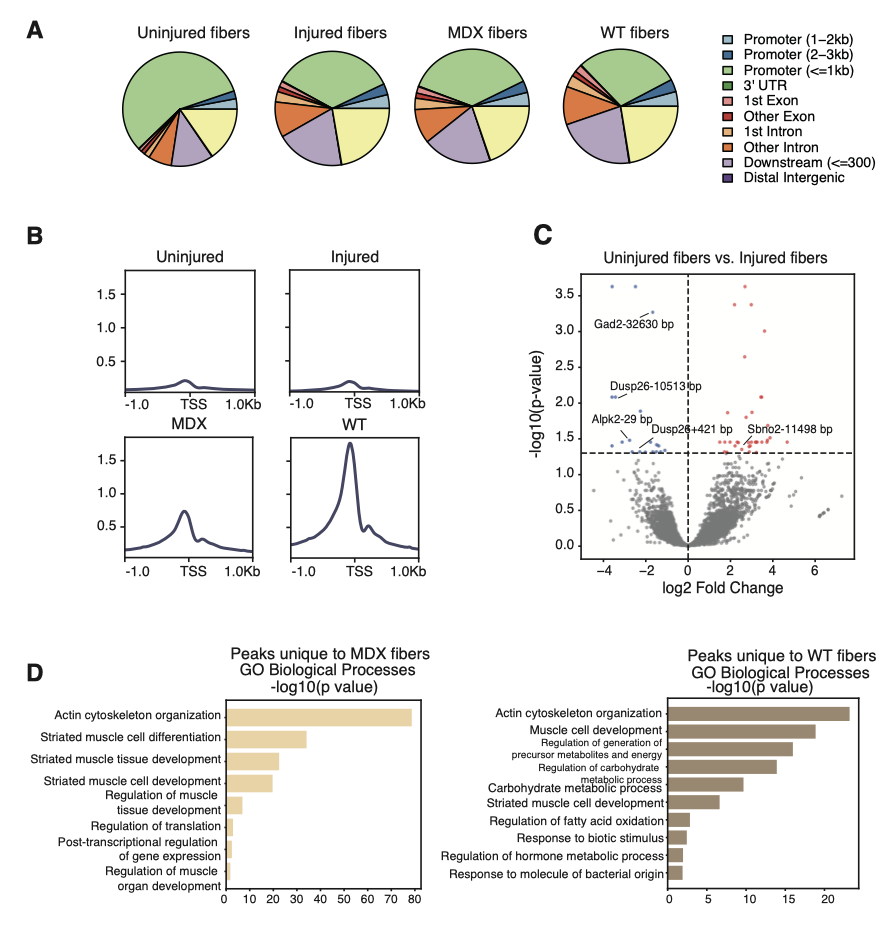

**Figure S4.** Reproduced biological patterns from a published ATAC-seq study. **A** Peak annotation pie charts for ATAC-Seq peaks of injured myofibers, uninjured myofibers, MDX myofibers and WT myofibers, respectively. **B** Enrichment of the signal at transcription start site (TSS) for the ATAC-Seq libraries of injured myofibers, uninjured myofibers, MDX myofibers and WT myofibers, respectively. **C** Volcano plot of differentially accessible peaks identified by FDR < 0.05 and LFC ≥ 1 between uninjured myofibers and injured myofibers. Each colored dot represents a differentially accessible peak and the distance to the nearest gene is annotated. **D** Gene Ontology (GO Biological Process) analysis of genes associated with unique peaks present in the MDX myofiber compared to WT myofibers and unique peaks present in the WT myofiber compared to MDX, respectively.
